# Deep nanoparticle protein corona plasma proteomics resolves a stage-specific peripheral signature of Alzheimer’s disease

**DOI:** 10.64898/2026.07.25.740710

**Authors:** Negar Mahmoudi, Bahareh Ghaffari, Kenneth Rogale, Samuel Cheeseman, Danilo Ritz, Alexander Schmidt, Liuchenxin Han, Qingling Li, Jared Deyarmin, Stephanie N. Samra, Dmitry Leshchiner, Sachi Horibata, Christopher Fowler, Borzoo Bonakdarpour, George Perry, Amir Ata Saei, Colin L. Masters, Babak Borhan, David Nisbet, Morteza Mahmoudi, the AIBL Research Group

## Abstract

**INTRODUCTION:** Alzheimer’s disease (AD) progresses over decades, yet plasma biomarkers that resolve disease stage rather than simply detect disease remain scarce. This distinction is clinically consequential because effective AD intervention depends on identifying patients before disease biology has progressed beyond a therapeutically responsive stage.

**METHODS:** We used small-molecule-modulated protein corona proteomics to profile plasma from 90 individuals in the Australian Imaging, Biomarker and Lifestyle cohort, stratified by Centiloid (CL) Aβ-amyloid burden (30 amyloid- negative, CL < 15; 30 moderate-to-high, CL 26 to 100; 30 very high, CL > 100). We quantified 3,176 proteins and applied differential abundance and actual causality analyses to identify stage-specific and candidate causal proteins.

**RESULTS:** Differential protein abundance was exclusively captured during the moderate-to-high AD transition, revealing a discrete proteomic “switch.” The switch was marked by accumulation of the autophagy receptor CALCOCO1, together with coordinated depletion of the S100A8/S100A9 calprotectin complex and core erythroid-cytoskeletal network structural markers (e.g., SPTA1, SPTB, ANK1). Adhesion G protein-coupled receptor G6 (ADGRG6) showed a significant moderate positive monotonic association with absolute CL burden, providing a proportional molecular anchor for cumulative disease burden. Actual causality analysis identified COL6A2, FOXRED2, P3H1, PRR4, and GOLGA5 as candidate upstream drivers linking matrix remodeling, Golgi trafficking, and collagen processing to AD progression.

**DISCUSSION:** These findings suggest a candidate blood-accessible framework for staging AD by active disease biology, which, if replicated in independent cohorts, may have implications for therapeutic selection and mechanism-guided clinical trials.

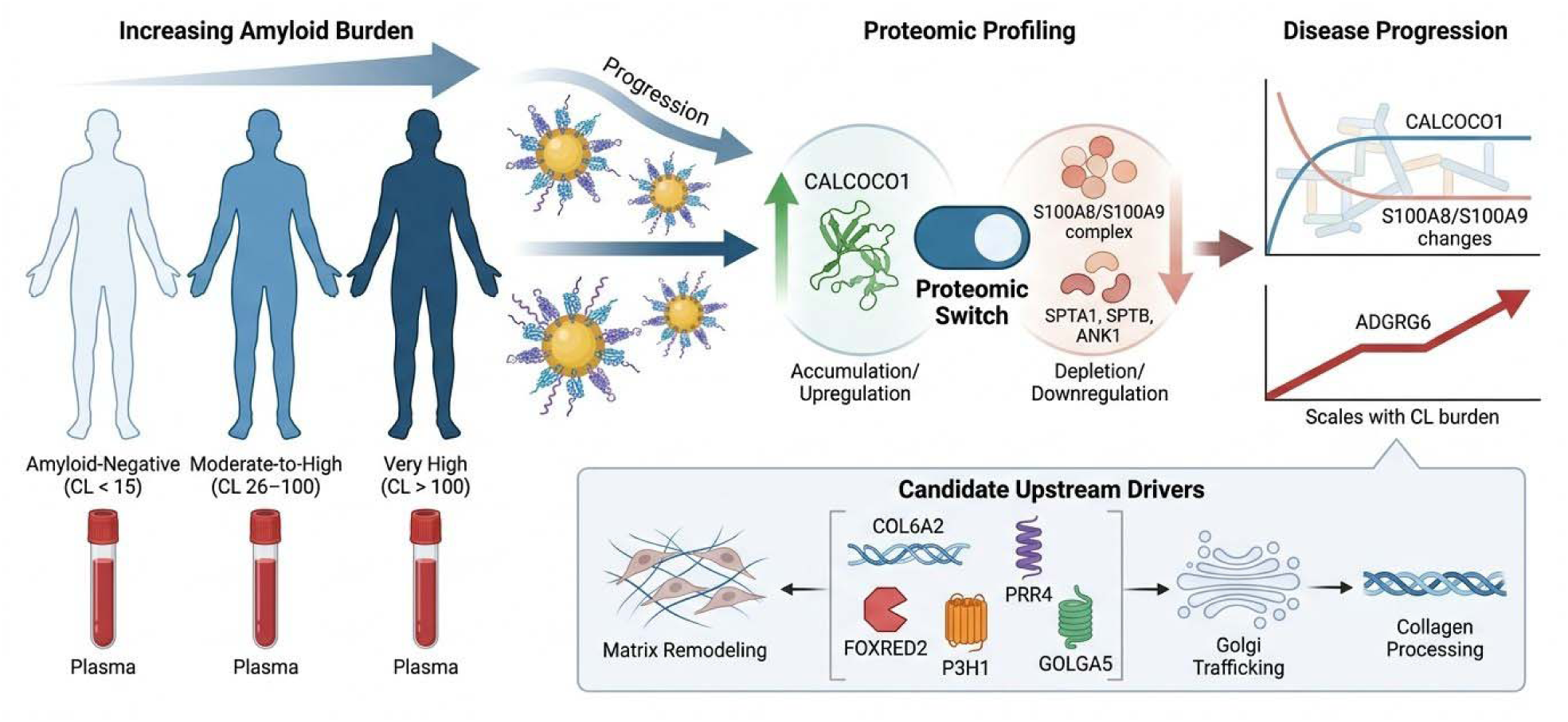

## Background

Alzheimer’s disease (AD) is a progressive neurodegenerative disorder characterized by the extracellular amyloid-β (Aβ) accumulation, intracellular tau aggregation, widespread synaptic dysfunction, and reactive cellular changes.^1–3^ A defining challenge in AD is that these disease processes unfold over decades, often beginning many years before measurable cognitive impairment.^2^ This prolonged preclinical phase has transformed AD from a disease defined primarily by dementia into a staged clinical and biological spectrum, in which the timing and dominance of specific molecular processes may determine whether a therapeutic intervention is likely to be effective.

The importance of biological staging has become increasingly urgent with the emergence of Aβ-directed therapies. Although Aβ burden remains central to AD diagnosis and patient selection, Aβ positivity alone does not define the active biological state of downstream reactive processes. Patients with similar Aβ burden may differ substantially in the extent of synaptic injury, tau accumulation, microglia and astrocyte activation, extracellular matrix (ECM) remodeling, and neuronal network collapse. This creates a therapeutic dilemma: an Aβ-directed intervention may be biologically appropriate during one stage of disease but insufficient, or poorly matched, once downstream degenerative programs have become dominant. Thus, the field needs blood-accessible biomarkers that do more than detect AD; it needs biomarkers that resolve disease stage, define the timing and extent of neuronal injury, and guide therapeutic strategy.

Current diagnostic advances highlight both the promise and limitations of blood-based biomarkers. Assays such as pTau217 (tau phosphorylated at threonine-217) can accurately detect AD changes in individuals with mild or even asymptomatic presentations.^4, 5^ Nonetheless, markers that report Aβ or tau-associated damage do not necessarily reveal when AD has transitioned from early molecular perturbation to broader neuronal network failure. This distinction may be critical for therapeutic decision-making and for clinical studies designed to evaluate mechanism-specific drugs.

Although genomics has advanced rapidly, the clinical application of proteomic data for AD has been hindered by the extreme complexity and dynamic range of the plasma proteome, together with the absence of protein amplification methods.^6–9^ Current workflows are often too cumbersome for large-scale clinical screening, limiting efforts to map the molecular landscape of neurodegeneration in blood. The plasma proteome contains more than 10,000 proteins spanning approximately 10 orders of magnitude in concentration, yet many disease-relevant proteins remain masked by high-abundance proteins such as albumin, immunoglobulins, and other carrier proteins.^10^

One approach for addressing plasma proteins complexity leverages the “protein corona”, the layer of biomolecules that spontaneously coats nanoparticles (NPs) when they are introduced to biological fluids.^11^ Protein corona formation can enrich a considerable portion of low-abundance proteins.^12, 13^ Nonetheless, unmodified corona formation is still influenced by abundant plasma proteins and does not fully overcome the masking effect of albumin and related carrier systems, nor does it capture the sub-proteome that is bound to albumin.^13^ To address this limitation, we recently developed a synergistic approach that combines NP protein corona formation with specific small- molecule modulators designed to bind albumin.^14, 15^ Rather than physically depleting albumin, these modulators alter albumin’s protein-binding dynamics, releasing associated passenger proteins into solution and increasing their availability for NP corona formation.^14^ At the same time, small molecule binding reduces affinity of albumin to the NP surface, creating physical space for the enrichment of these newly released, disease-relevant proteins.^14^ We reasoned that this dual strategy would enrich low-abundance proteins and extend the depth of plasma profiling achievable across the AD continuum.

Most contemporary plasma proteomics studies rely heavily on AI and machine learning approaches that identify disease-associated patterns through correlation, but not necessarily causation.^16–18^ To further enhance our capability of identifying causal plasma proteins with a strong association with AD, we incorporated the concept of “actual causality”, as formalized by Halpern and Pearl,^19, 20^ into our analytical pipeline. This framework is designed to move beyond correlation by identifying proteins whose variance is more consistent with direct involvement in disease-state transitions or causal indication of disease-related changes.

## Methods

### Plasma samples

Plasma samples from 90 participants were chosen from the Alzheimer’s disease (AD) cohort of the AIBL Study. Among them, 30 individuals were negative for AD, and 60 were positive with varying Aβ Centiloid scales. This longitudinal cohort, established in 2006, tracks thousands of Australians to study AD progression, pioneering early diagnosis through Aβ biomarkers and blood tests.^21^

### Standardized Aβ and Tau PET quantification

A central feature of this cohort’s clinical validation is the use of the Centiloid (CL) scale, which provides a standardized framework to harmonize Aβ PET imaging across different tracers and scanners.^22^ This scale anchors a typical Aβ burden at 0 for young controls and 100 for patients with mild AD. To improve longitudinal reliability when tracers or scanners are switched, a novel non-negative matrix factorization (NMF) Centiloid method is employed.^23^ Based on these values, Aβ status is categorized into various risk levels: Negative (<15 CL), Uncertain (15–25 CL), Moderate-High (26-100 CL), and Very High (>100 CL). Notably, the risk of progression to dementia increases significantly with these burdens, rising from 5% in the negative/uncertain groups to 50% in the very high category. Tau deposition is similarly staged using a stereospecific approach to track the spread of tau from the mesial-temporal lobe (Me) into the temporoparietal (Te) and meta-temporal (MT) neocortical regions. These categorical measures are essential for disease staging and prognosis, as both the amount and anatomical location of tau deposits are highly relevant to clinical progression.

### Structural MRI and white matter analysis

The clinical characterization of these plasma samples is further supported by rigorous MRI volumetric analysis using the CurAIBL platform.^24^ This includes standardized estimates for hippocampal, cortical grey matter, and ventricular volumes, all of which are corrected for head size (intracranial volume) and scanner-specific systematic errors to ensure cross-subject comparability. These structural metrics are critical for identifying disease-related atrophy, particularly in the hippocampus, which is defined using the “Harmonized Hippocampus Protocol”. Beyond gray matter atrophy, the cohort is assessed for White Matter Hyperintensities (WMH) using neural network classifiers to quantify total, periventricular, and deep WMH volumes.^25^

### Small molecule protein corona

The platform contains polystyrene nanoparticles with the size of 200 nm and phosphatidylcholine as a small molecule. Full characterization details of the nanoparticles’ protein corona platform and its optimization for achieving pure protein corona without protein aggregates are available in our recent publications.^14, 15, 26^

### Liquid chromatography-tandem mass spectrometry (LC-MS/MS)

Dried peptides were reconstituted in 0.1% aqueous formic acid (FA) supplemented with 0.02% n-dodecyl-β-D-maltoside (DDM) to maximize solubility and minimize adsorptive loss. Chromatographic separation was performed on a Vanquish Neo UHPLC system (Thermo Fisher Scientific) coupled to a timsTOF Ultra 2 mass spectrometer (Bruker) via a CaptiveSpray nano-electrospray source. Peptides were resolved on a 30 cm in-house packed analytical column (100 µm I.D., ReproSil Saphir 100 C18, 1.5 µm; Dr. Maisch GmbH) maintained at 60°C. Separation was executed at a flow rate of 400 nL/min using a binary gradient: (Buffer A: 0.1% FA in water; Buffer B: 80% ACN, 0.1% FA in water). The 35-minute gradient progressed from 2% to 25% B over 25 minutes, then to 35% B over 5 minutes. The column was washed at 95% B for 4 minutes and re-equilibrated at 2% B.

MS data were acquired in dia-PASEF mode (cycle time ∼0.95 s) over a mass range of 100–1700 m/z. The dia-PASEF scheme utilized 8 scans with 3 ion mobility windows per scan, targeting a precursor range of 400–1000 m/z with 25 Da isolation windows. Ion mobility (1/K_0_) was set from 0.64 to 1.37 V·s/cm². Collision energy was linearly ramped from 20 eV at 1/K_0_ = 0.6 to 59 eV at 1/K_0_ = 1.6. Source parameters included a capillary voltage of 1600 V and a dry gas flow of 3 L/min at 200°C, with Ion Charge Control (ICC 2.0) enabled.

### Data processing and bioinformatic analysis

Raw files were processed in Spectronaut (v19.0, Biognosys) using the directDIA+ workflow. Spectra were searched against the UniProt *Homo sapiens* reference proteome (20,360 sequences, Feb 2022) and 392 common contaminants. Cross-Run Normalization was disabled to preserve raw quantitative integrity. Quantitative results were exported via the Pivot Report function for statistical analysis. Data analysis and visualization were conducted using custom Python scripts and GraphPad Prism (v10.6.1). Samples were analyzed in triplicate. Prior to quantification, known contaminants (prefix “Con–”) were removed. Missing values were handled as follows: i) single missing replicate (imputed using the mean of the remaining two replicates; and ii) double missing replicates (the protein was excluded for that specific sample. Protein intensities were normalized to the total protein intensity per replicate and expressed as relative abundance percentages. This compositional approach ensured robust comparisons across the different experimental conditions. Differential expression was assessed using the limma package^27^ with a no-intercept design matrix encoding three groups: Negative, Moderate-High, and Very High. Raw protein abundance values were loaded from a pre-normalized, contaminant-filtered dataset. Proteins were retained if they were detected in at least 50% of subjects within at least one group, thereby reducing the influence of highly sparse features. Remaining missing values were imputed with a small constant (1 × 10⁻) to represent non-detected measurements as very low-abundance signals, allowing the analysis to capture differences associated with both reduced measured abundance and increased non-detection. Retained abundances were log₂-transformed. Three pairwise contrasts were tested: Moderate-High vs. Negative, Very High vs. Negative, and Very High vs. Moderate-High. Significance was evaluated using both nominal p-value (<0.05) and Benjamini–Hochberg-adjusted *p*-value (FDR <0.1), with a |log₂FC| threshold of 0.25 for volcano plot annotation due to data sparsity.

### Two-species plasma dilution series for quantitative accuracy validation

Plasma digest samples were analyzed using a Thermo Scientific Vanquish Neo UHPLC system operated in a trap-and-elute configuration and coupled to a Thermo Scientific Orbitrap Astral Zoom mass spectrometer. For the 24 samples-per-day (SPD) workflow, peptides were separated on an EASY-Spray HPLC ES7550PN column (75 μm × 50 cm, 2 μm particle size) using a 54-minute chromatographic gradient. For the 60 SPD workflow, analytical separation was performed on an EASY-Spray PepMap HPLC ES906 column (150 μm × 15 cm, 2 μm particle size) with a 22.35-minute gradient. Detailed liquid chromatography gradient conditions for the 24 SPD and 60 SPD workflows are provided in Supplemental Table 1(A) and 1(B), respectively. For each analysis, 500 ng of peptides from individual plasma sample were injected onto the analytical column. Eluting peptides were analyzed by narrow-window data-independent acquisition (nDIA) on the Orbitrap Astral Zoom mass spectrometer. The acquisition parameters for the full-scan MS1 and nDIA MS2 experiments are provided in Supplemental Tables 2(A)-(C) for 24- and 60-SPD methods. For the human–chicken plasma dilution series, each sample preparation replicate containing the full set of dilution levels was analyzed as a separate block, with blank injections inserted between replicate blocks. Within each replicate block, samples were injected in a randomized order with the constraints that the 99% chicken plasma/1% human plasma (99C/1H) sample was not adjacent to the 0C/100H or 30C/70H samples, and the 95C/5H sample was not adjacent to the 0C/100H sample. These constraints were applied to minimize transient ion suppression, retention time shifts and carryover-like artifacts caused by abrupt changes in sample matrix composition. Analytical quality control was performed by analyzing the same plasma sample before and after the analytical sequence. Linearity, precision and accuracy were assessed using the dilution series and sample preparation replicates. ***Data processing.*** All mass spectrometry raw data were analyzed using Spectronaut v20 (Biognosys AG) in DirectDIA mode with a library-free workflow. Human (UniProt Organism ID: 9606; 20,421 entries, Swiss-Prot–reviewed canonical sequences) and chicken (UniProt Proteome ID:9031; 43,712 entries) protein FASTA files were downloaded from UniProt and used for the library-free database searches. DirectDIA searches were performed using the default Biognosys (BGS) factory settings. More specifically, enzyme and modification parameters were set at trypsin enzyme specificity with a maximum of two missed cleavages, carbamidomethylation of cysteines (fixed modification), and oxidation of methionine and N-acetylation of proteins (as variable modifications). Spectral matching of the peptides was performed with a FDR of 1% identified proteins with a minimum of two peptides for protein identification.

### Data availability

Proteomic data have been deposited to the ProteomeXchange Consortium (https://www.proteomexchange.org/) via the MassIVE partner repository (https://massive.ucsd.edu/) with MassIVE data set identifier MSV000102244 and ProteomeXchange identifier PXD080029.

## Results

In this study, we employ small-molecule-modulated protein corona proteomics to a clinically characterized cohort spanning the AD continuum. Our objective was not simply to discover additional AD-associated plasma proteins, but to determine whether deep plasma proteomics can resolve stage-specific disease status and identify molecular transitions with therapeutic significance. To move beyond mere statistical association, we implemented actual causality to mathematically identify specific plasma proteins that drive or directly indicate AD. Our study identifies increased CALCOCO1 abundance, together with coordinated systemic decreases in the abundance of the S100A8/S100A9 glial reactive complex and core erythroid-cytoskeletal structural markers, including SPTA1, SPTB, and ANK1, as high-performance candidate blood-based biomarkers associated with AD progression. We further identified ADGRG6 as the only protein showing a positive correlation with Centiloid (CL) levels. Together, these findings support a stage-resolved model of AD in which deep plasma proteomics can inform two critical decisions; when therapeutic intervention may be appropriate, and which molecular features provide the strongest foundation for translating proteomic discoveries into clinical diagnostic tools.

### Small-molecule corona proteomics profiles Aβ-staged AD plasma

AD therapy suffers from variable efficacy across patient cohorts in part because patients are often staged by disease burden rather than by the active molecular state of the disease. To determine whether blood-based proteomics could resolve this question, we applied our small molecule (phosphatidylcholine) modulated protein corona platform^14, 15^ to 90 plasma samples obtained from the Australian Imaging, Biomarker, and Lifestyle (AIBL) Study, a longitudinal cohort established in 2006 to investigate the transition from healthy aging to AD.^21^ The cohort included 30 amyloid-negative controls (mean age 74 ± 6 years) and 60 amyloid-positive individuals, spanning a broad range of Aβ accumulation (mean age 74 ± 7 years), quantified using the Centiloid (CL) scale (see the Method section for details).^23^ Aβ-negative status was defined as CL <15, whereas Aβ-positive individuals were stratified into moderate-high amyloid burden (26–100 CL) and very high amyloid burden (>100 CL) according to Non-Negative Matrix Factorization (NMF) CL-based classification.^28–30^ This cohort design allowed us to ask whether plasma proteomic changes scale gradually with Aβ accumulation or instead reveal discrete biological transitions that could inform therapeutic staging.

Sample selection incorporated multi-modal clinical data, including standardized hippocampal volumetric estimates obtained via CurAIBL and Tau-PET imaging to stage neurodegenerative progression from the mesial-temporal lobe to the neocortex.^24^ Thus, proteomic analyses were anchored to established imaging and biomarker features of AD, rather than to clinical diagnosis alone. Mass spectrometry analysis of the small-molecule-modulated protein corona identified 3,176 distinct proteins, with 2,894 proteins shared across both AD-negative and AD-positive groups (**Figure 1**). This overlap provided a consistent baseline for detecting shifts in protein abundance as a function of Aβ burden and disease stage.

**Figure 1.**
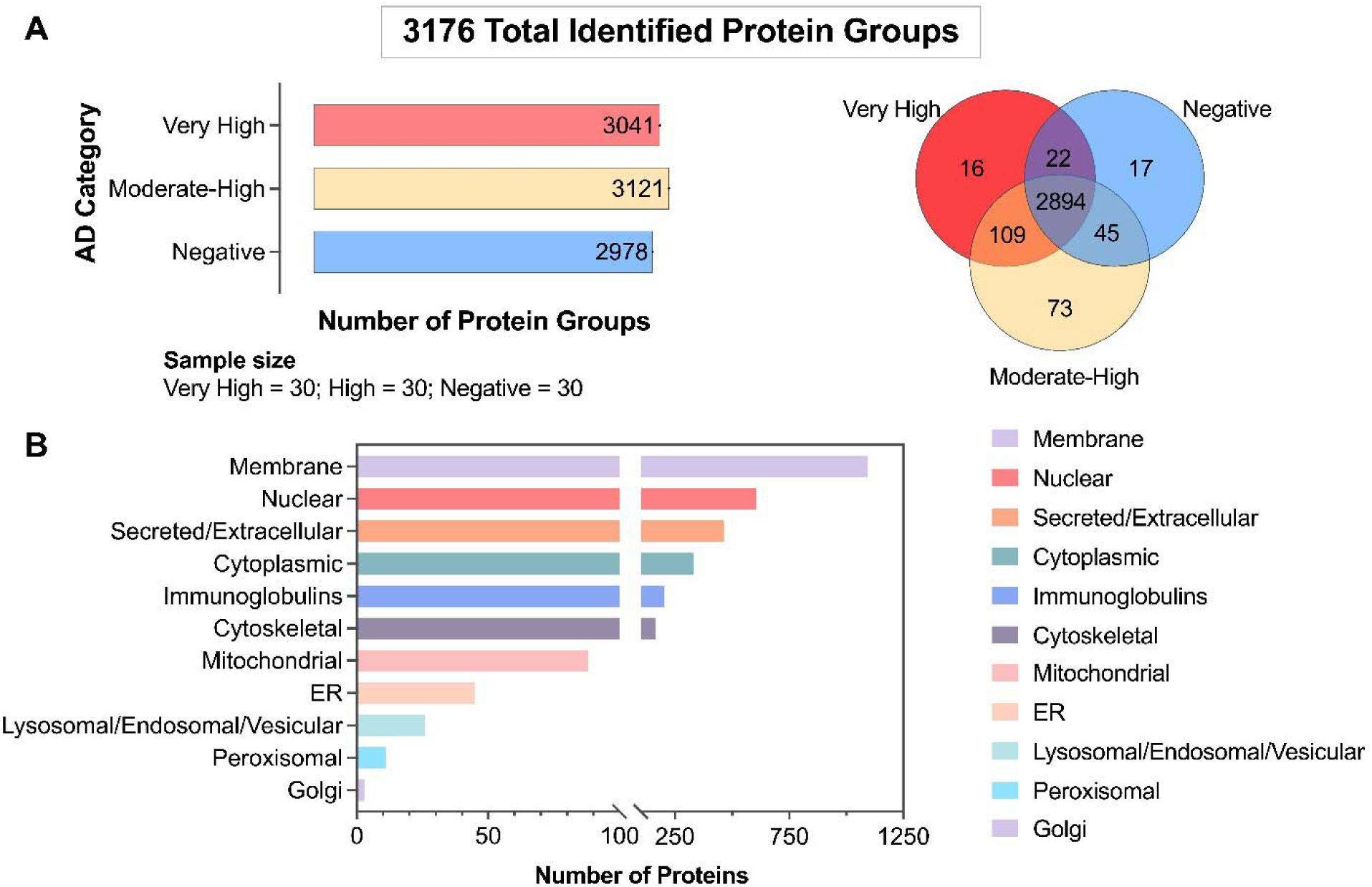
Deep plasma proteomic profiling and cohort stratification using small-molecule-modulated protein corona. **A)** Analysis of 90 plasma samples from the AIBL longitudinal cohort categorized by Aβ Centiloid (CL) scale: Negative (<15 CL), Moderate-High (26–100 CL), and Very High (>100 CL). The Venn diagram illustrates the distribution of the 3,176 total identified protein groups, showing 2,894 proteins shared across all clinical categories. **B)** Functional and subcellular classification of the 3,176 identified protein groups based on protein annotation. Proteins were categorized as membrane, nuclear, secreted/extracellular, cytoplasmic, immunoglobulin, cytoskeletal, mitochondrial, endoplasmic reticulum (ER), lysosomal/endosomal/vesicular, peroxisomal, or Golgi-associated. The x-axis indicates the number of proteins assigned to each category and is discontinuous to accommodate the wide range in category sizes.

Although our small-molecule-modulated protein corona platform has achieved substantially deeper proteomic coverage, identifying over 6,000 unique protein groups in freshly isolated plasma,^13, 14^ the number of quantified proteins in historical biobanked cohorts, such as the AIBL longitudinal cohort, is appreciably lower. This difference likely reflects the physical nature of corona formation, which can capture circulating extracellular vesicles (EVs) together with soluble plasma proteins and thereby increase apparent discovery depth.^31^ The plasma-resident EVs in long-termed biobanked samples, however, are susceptible to structural degradation and physical disruption during long-term ultra-low temperature storage, including damage associated with ice-crystal formation, with additional vulnerability introduced by repeated freeze-thaw cycles.^32–35^ Cryopreservation-induced membrane lysis and exosomal degradation can reduce intact vesicular structures, surface antigens, and trapped cargo, thereby decreasing the pool of broad-spectrum, low-abundance proteins available for nanoparticle capture relative to freshly processed plasma matrices.

Having established a clinically anchored, Aβ-staged plasma proteomic dataset, we next asked whether the peripheral proteome reports AD progression as a gradual continuum, a discrete biological transition, or both. The results therefore proceed in three steps: first, we use differential abundance analysis and analyze volcano plots to define stage-specific proteomic phenotypes across the CL continuum; second, we apply actual- causality analysis to prioritize protein-state assignments that behave as candidate upstream drivers under matched counterfactual comparisons; and third, we integrate these two layers to connect blood-detectable staging markers with the biological processes that may determine therapeutic state.

To confirm that the differential protein abundances observed across the Aβ-staged AIBL cohort reflect genuine, quantitatively accurate changes in protein levels^36^ rather than platform-specific noise or non-linear detection artifacts, we performed an orthogonal validation experiment using a two-species plasma dilution series (**Figure 2A**). Human plasma was mixed with chicken plasma across a defined series (1, 5, 10, 20, 40, 60, 80, and 100% human plasma by volume) and analyzed by DIA-MS on two independent platforms at high throughput (24 and 60 samples per day (SPD); **Supplemental Tables 1-3**). Because the human and chicken plasma proteomes are sufficiently divergent, human-specific peptides and proteins detected at each dilution step provide a ground-truth benchmark against which measured signal intensities can be directly compared to their known, expected proportions (**Figure 2B**). The cross-species plasma analysis highlighted the depth of proteome coverage achieved by both methods, with SPD24 quantifying 6,658 protein groups and 93,806 peptides and SPD60 quantifying 5,866 protein groups and 76,473 peptides. The large majority of features (84.4–85.8% of proteins; 69.3–72.6% of peptides) showed a coefficient of determination (R²) of at least 0.90 between measured normalized signal intensity and expected human plasma fraction, with median R² values of 0.97–0.98 for proteins and 0.95–0.96 for peptides (**Figure 2C**). Individual proteins spanning a wide abundance range and relevant to this study’s biological findings — including coagulation factors (F10, F5), the acute-phase protein LRG1, and lower-abundance regulatory and translational proteins (GSPT1, RPS8, EIF2A, ANKS1A, GARS1, MCTS1, VWA5A) — each tracked their expected dilution fraction with R² values exceeding 0.99 (**Figure 2D**). These results demonstrate that our small-molecule-modulated protein corona platform preserves a linear, quantitatively accurate relationship between true protein abundance and measured signal across a greater than 100-fold concentration range and across proteins of widely differing plasma abundance. This orthogonal validation supports the conclusion that the stage-specific abundance changes reported for AD-associated proteins in this study reflect real, proportionally scaled biological differences in circulating protein levels rather than technical artifacts of the corona-capture or mass spectrometry workflow.

**Figure 2.**
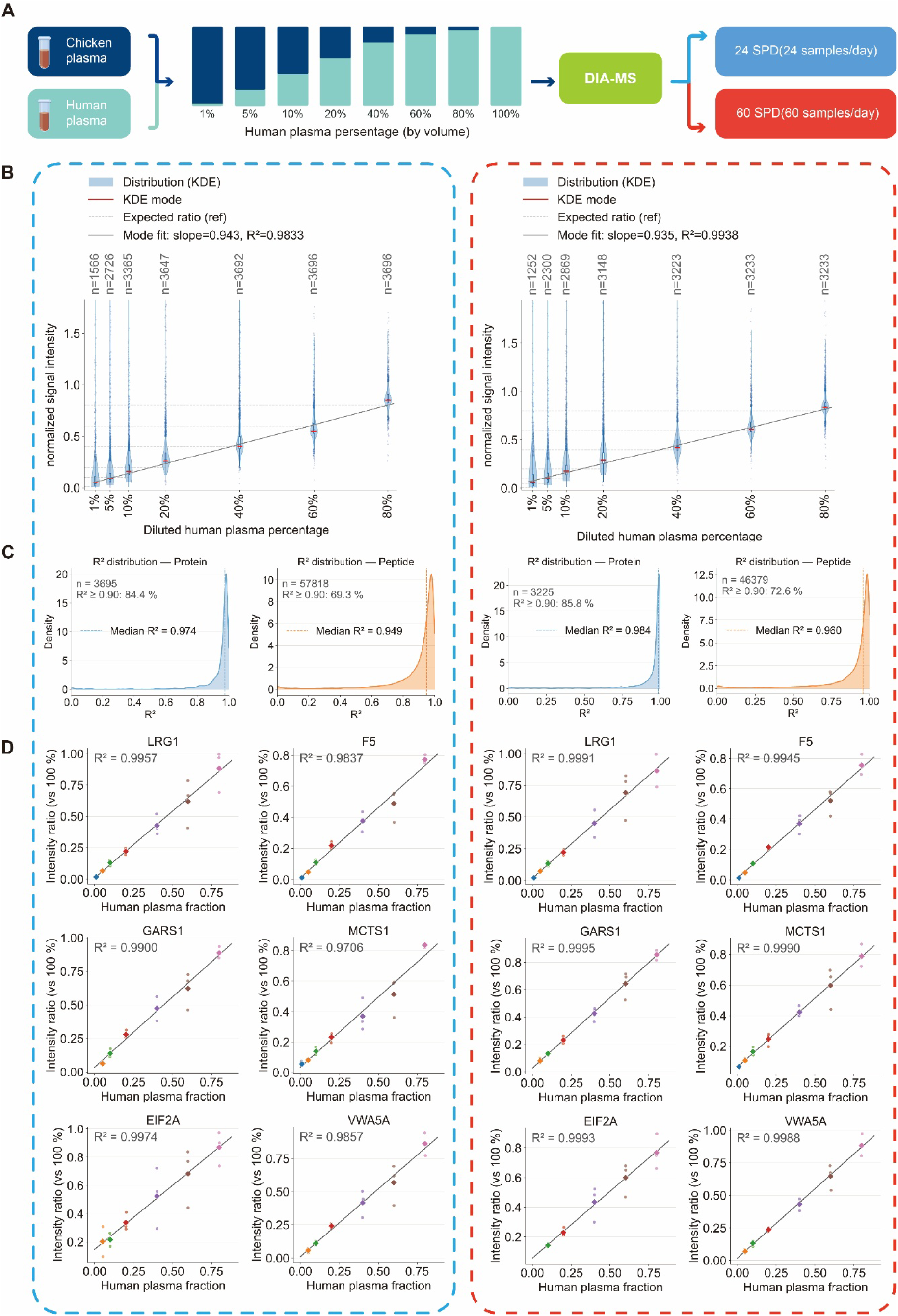
Dual-acquisition evaluation of protein-level quantitative accuracy in a human–chicken plasma dilution series. **A)** Overview of the human–chicken plasma dilution design (1–100% human plasma) and DIA-MS workflow, analyzed at two acquisition throughputs, SPD24 and SPD60. **B)** Quantitative linearity of protein intensity ratios (relative to the 100% human plasma reference) across the dilution series, for SPD24 (left) and SPD60 (right). Violins show the per-group distribution; red bars, KDE mode; grey dashed lines, theoretical expected ratio; black line, linear fit through group modes (slope, R² annotated). n, proteins quantified per group. **C)** Distribution of per-protein R² (observed ratio vs. nominal human fraction, OLS regression) for SPD24 (left) and SPD60 (right). Dashed line, median R²; inset, n and percentage with R² ≥ 0.90. **D)** Representative high-linearity proteins across high-, low-, and ultra-low-abundance tiers (HPA-defined; one tier per row, two proteins per tier), for SPD24 (left) and SPD60 (right). Points, replicate measurements; diamonds, replicate means; line, OLS fit (R² annotated). Proteins shown differ between instruments (selected independently).

### Stage-specific differential expression reveals a high-impact peripheral “switch”

We performed pairwise differential expression analysis across the Aβ-stratified cohorts (**Figure 3A-D**). Differential protein expression, defined by adjusted *p* < 0.1, was captured only during the moderate-to-high AD transition (Moderate-High *vs.* Negative; (**Figure 3A** and **B**, and **Figure 4A** and **B**)). In contrast, comparisons involving the advanced stage, Very High vs. Negative and Very High vs. High, did not yield proteins that cleared the adjusted significance threshold (**Figure 3C** and **D**). These results are consistent with the moderate-to-high transition representing a coordinated and relatively specific peripheral molecular “switch,” although we cannot exclude that reduced sample integrity in longer-stored Very High specimens contributed to the absence of further significant changes at higher CL.

**Figure 3.**
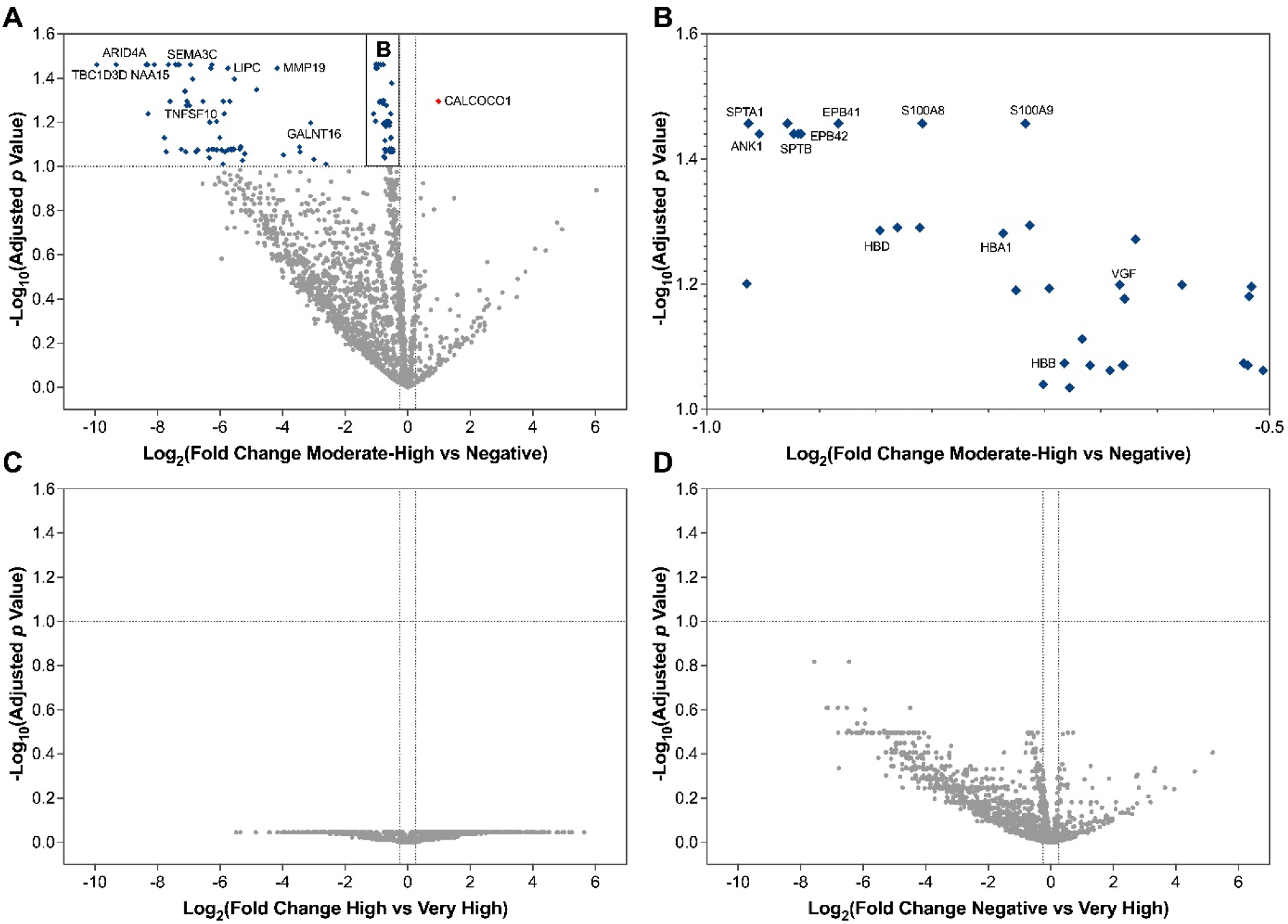
Stage-specific plasma proteomic alterations across Aβ burden groups. **A)** Volcano plot showing pairwise differential abundance analysis of the quantified plasma proteins between the moderate-to-high Aβ burden cohort, defined as 26 ≤ CL < 100, n = 30, and the Aβ-negative control cohort, defined as CL < 15, n = 30. Each point represents one protein, with log_₂_ fold change plotted on the x-axis and −log_₁₀_ FDR-adjusted *p* value plotted on the y-axis. The horizontal dashed line indicates the significance threshold of adjusted *p* < 0.1, and the vertical dashed lines indicate the applied log_₂_ fold- change cutoff. Significant proteins enriched in the moderate-to-high cohort are shown on the right, including CALCOCO1, while significant proteins depleted in the moderate-to-high cohort are shown on the left. These depleted proteins include immune-related proteins such as the S100A8/S100A9 calprotectin complex, erythroid and cytoskeletal markers including SPTA1, SPTB, ANK1, EPB41, and EPB42, hemoglobin-associated proteins including HBA1, HBB, and HBD, and members of the TBC1D3 family. Non-significant proteins are shown in gray. **B)** Magnified view of the significant depleted region from panel A, highlighting selected proteins within the structural, immune-related, and hemoglobin-associated clusters. **C)** Moderate-to-high versus very-high Aβ burden comparison, where the very-high cohort is defined as CL > 100, n = 30, showing no proteins passing the adjusted significance threshold. **D)** Aβ-negative versus very-high Aβ burden comparison, showing broader abundance variation but no proteins meeting the adjusted significance cutoff under the applied criteria.

**Figure 4.**
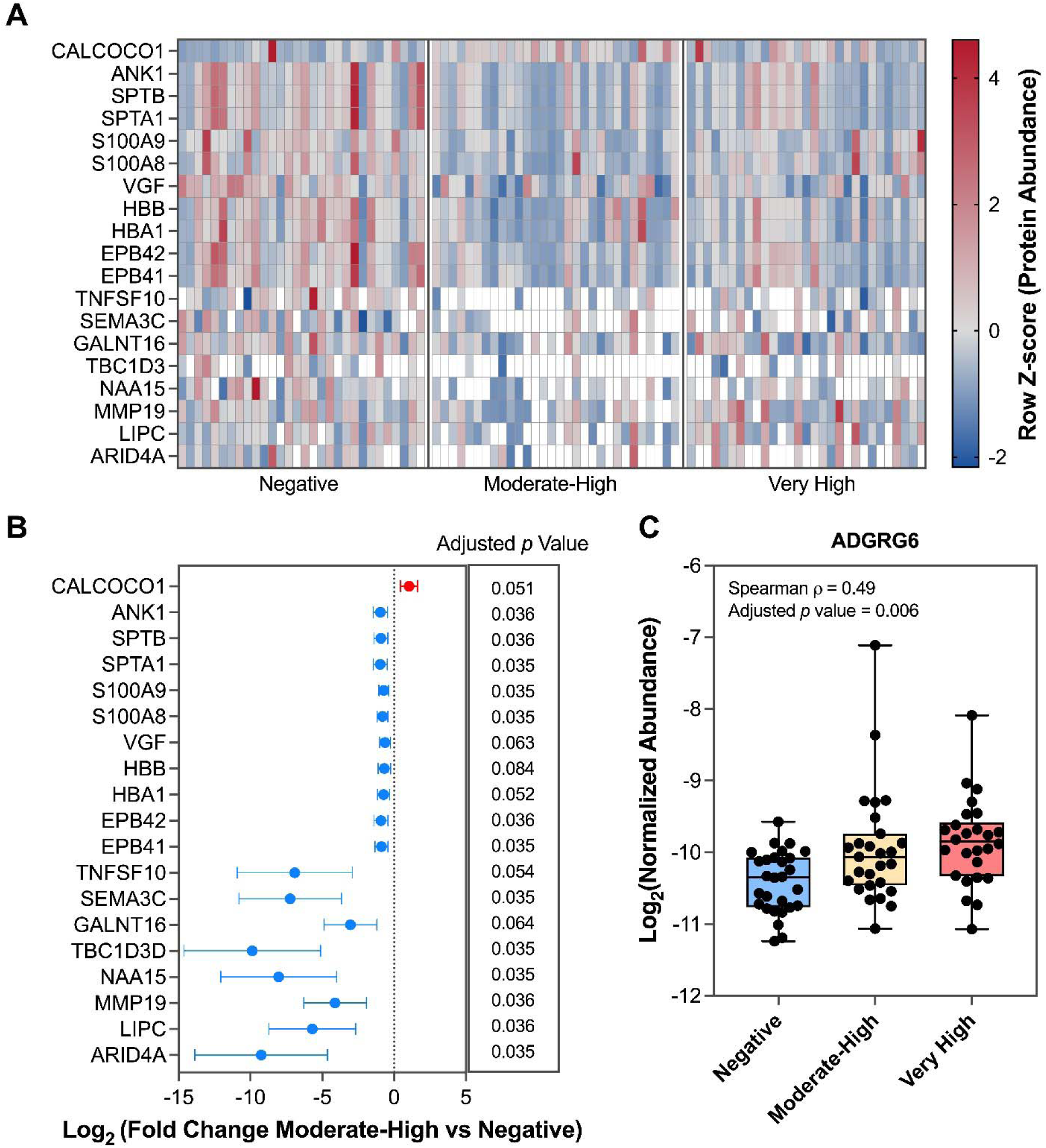
Differential plasma protein abundance across Alzheimer’s disease amyloid-burden groups. **A)** Heatmap showing the relative abundance of selected differentially abundant proteins across Negative, Moderate–High, and Very High amyloid groups. Abundance values were standardized by protein using row Z-scores, with white cells indicating non-detected measurements. Each column represents an individual subject, and subjects are grouped according to amyloid category. **B)** Forest plot showing log_₂_ fold changes for selected proteins in the Moderate–High group relative to the Negative group. Points indicate estimated log_₂_ fold changes, horizontal error bars represent 95% confidence intervals, and the dashed vertical line denotes no change (log_₂_ fold change = 0). Corresponding FDR- adjusted *p* values are shown on the right. **C)** Distribution of log_₂_-normalized ADGRG6 abundance across the three amyloid groups, with individual subject values overlaid. ADGRG6 abundance showed a moderate positive association with continuous CL values across the full clinical cohort (Spearman’s ρ = 0.49, adjusted *p* = 0.006).

On the positive fold-change side of the volcano plot, the most prominent change was the accumulation of CALCOCO1 (Calcium-binding and coiled-coil domain-containing protein 1). CALCOCO1 is a principal intracellular cargo receptor for reticulophagy, mediating the targeted autophagic clearance of endoplasmic reticulum (ER) sub- compartments.^37–39^ Its significant increase in abundance may reflect an acute compensatory response aimed at clearing accumulated misfolded proteins and ER- stress–associated aggregates during early-stage AD.

The direction of this change merits comment, because CALCOCO1 is a soluble, non-secreted intracellular autophagy receptor, and its presence in plasma at all is most parsimoniously explained by packaging into circulating EVs rather than by conventional secretion. Our small-molecule-modulated protein corona platform is known to co-enrich EV-associated proteins alongside soluble plasma proteins^31^, which is the most likely route by which an ER/Golgi-resident receptor such as CALCOCO1 becomes detectable in this dataset at all. This is consistent with independent reports that blood-derived EVs and particles carry brain-derived, disease-discriminating RNA and protein cargo detectable non-invasively, and that a protein corona can form on the surface of such particles in blood plasma.^40^ Mechanistically, CALCOCO1 is itself degraded together with its ER and Golgi cargo during successful reticulophagy and Golgiphagy^37–39^; loss-of-function experiments have shown that when the downstream machinery required for this turnover is limiting (e.g., depletion of the VAP proteins that CALCOCO1 requires for cargo engagement), CALCOCO1 itself accumulates rather than being cleared^38^, in the same way that other selective-autophagy receptors (e.g., SQSTM1/p62) accumulate when autophagic-lysosomal flux is outpaced by demand.^41^ We propose that a similar imbalance — increased CALCOCO1-mediated reticulophagy/Golgiphagy demand during the moderate-to-high transition, combined with limited downstream clearance capacity — could plausibly drive both intracellular CALCOCO1 accumulation and, via increased EV-mediated release from stressed cells^42^, the elevated EV-associated CALCOCO1 we observe in plasma. This would be consistent with our finding that the CALCOCO1 increase is confined to the moderate-to-high transition and returned to its negative level at very high amyloid burden, if the compensatory response itself becomes exhausted, or its cellular source is progressively lost, at later disease stages. We note that a newly published, large-scale plasma proteomic study of neurodegenerative dementias also identified CALCOCO1 among the strongest AD-specific markers, but reported it as progressively downregulated across the AD continuum, consistent with an earlier multi-omics report.^43, 44^ This finding directly opposes the CALCOCO1 accumulation we describe here and warrants explicit comment. Both prior reports quantified CALCOCO1 using large-scale antibody-based platforms (proximity extension assay and related affinity methods) applied to neat, unfractionated plasma, in which detection depends on epitope accessibility to solution-phase antibody binding. Because CALCOCO1 lacks a signal peptide and is not conventionally secreted, virtually all circulating CALCOCO1 must arise through a non-canonical route, most plausibly packaging into EVs released from autophagically stressed cells, rather than free secretion into plasma. Antibody-based detection of intracellular, EV-packaged cargo in biofluids is known to be strongly method- and platform-dependent, varying with vesicle integrity and isolation protocol.^45, 46^ Our small-molecule-modulated protein corona platform co-enriches EV-associated material alongside soluble plasma proteins and subsequently denatures and digests the entire captured fraction for mass spectrometry, and would therefore be expected to report a pool of CALCOCO1 inaccessible to solution-phase immunoassays.^47^

To test this possibility directly, we compared CALCOCO1 recovery in the nanoparticle protein corona formed on neat human plasma versus the same plasma systematically depleted of EVs by immunoaffinity capture, using our previously validated EV-depletion platform^47^. CALCOCO1 was consistently detected in the corona formed on neat plasma across all six nanoparticle sizes tested (50–1000 nm), whereas in EV-depleted plasma it was undetectable at most nanoparticle sizes and markedly reduced at the remainder relative to their matched neat-plasma conditions (**Figure 5**). Because EV depletion removes the vesicular compartment while leaving the soluble plasma protein pool intact, this near-complete loss of detectable CALCOCO1 upon EV depletion indicates that the CALCOCO1 signal recovered by our platform originates predominantly from EV-packaged, rather than free, soluble protein. This result provides direct empirical support for a platform-dependent, compartmental-sampling explanation for the discrepant CALCOCO1 findings across studies: CALCOCO1 behaves as an EV-cargo protein in plasma, and its detectability is therefore governed largely by whether a given analytical platform captures the vesicular fraction, as our nanoparticle protein corona approach does, or is instead restricted, by design or by antibody accessibility, to the soluble, free-protein fraction, as is the case for solution-phase immunoassays such as proximity extension assay. This finding also raises a broader methodological caution: for any plasma protein whose abundance is substantially EV-derived, the analytical platform used, and not only underlying disease biology, may determine whether the protein is detected at all, and even the apparent direction of a disease-associated change, when findings are compared across proteomic platforms with differing access to the vesicular compartment.

**Figure 5.**
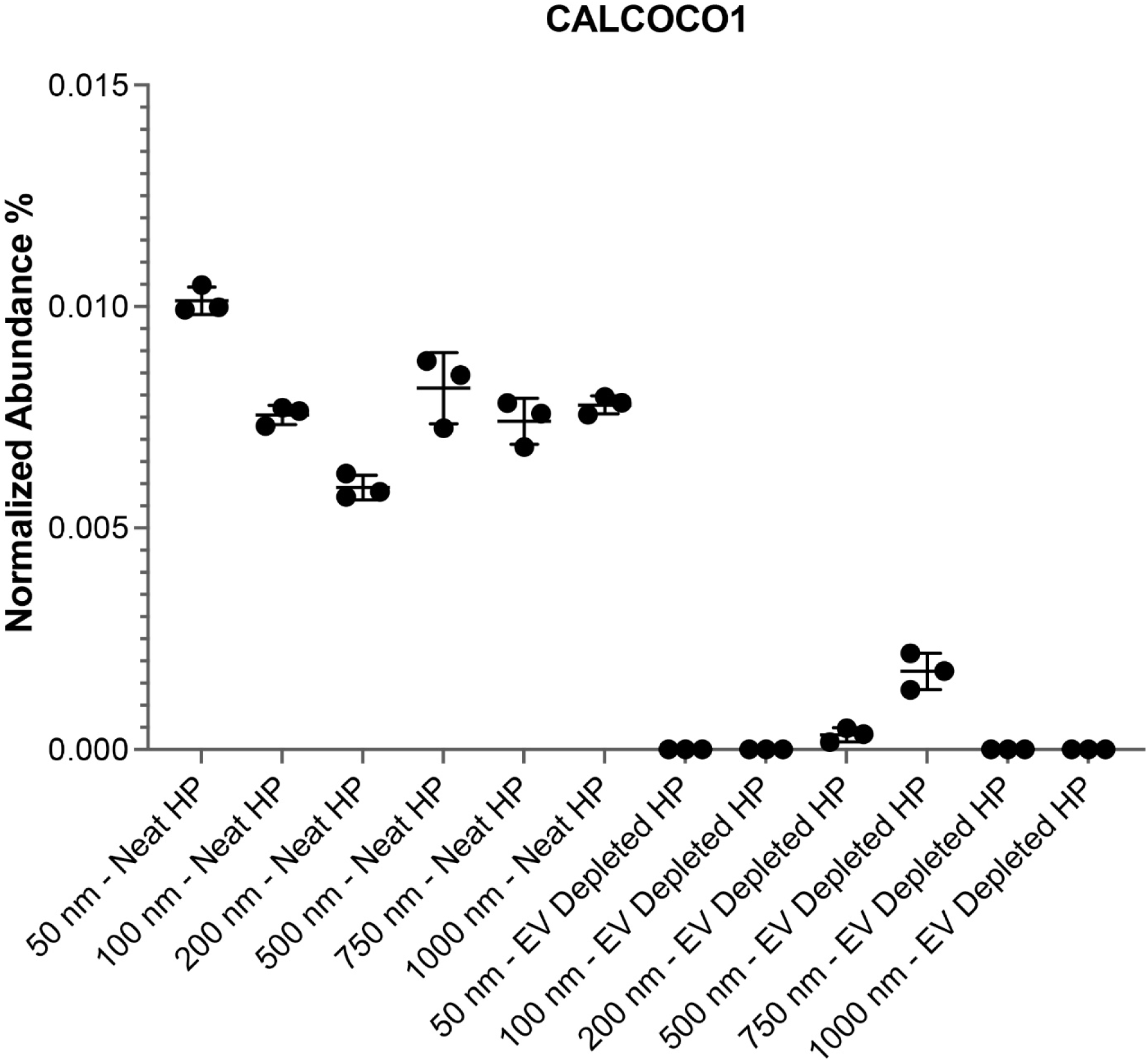
EV depletion abolishes corona detection of CALCOCO1, confirming its extracellular- vesicle origin. Normalized abundance (%) of CALCOCO1 in the protein corona formed on nanoparticles ranging from 50 to 1000 nm in diameter, incubated with either neat human plasma or the same plasma systematically depleted of EVs by immunoaffinity capture. Individual points represent three technical LC– MS/MS replicates per condition; error bars indicate mean ± SD; values plotted at zero indicate that CALCOCO1 was not detected. CALCOCO1 was consistently detected across all nanoparticle sizes tested in neat plasma but was absent or markedly reduced in most EV-depleted plasma conditions.

An important caveat to this compartmental interpretation is that EVs are not perfectly membrane-intact throughout the pre-analytical pipeline: delayed processing, repeated freeze–thaw cycling, and long-term storage are well documented to compromise EV membrane integrity and release luminal cargo into the surrounding fluid.^32–35, 48^ Because CALCOCO1 itself lacks a transmembrane domain and a canonical secretory signal peptide, any luminal CALCOCO1 released by such ex vivo vesicle disruption, whether occurring in the circulation before collection or during downstream sample handling, would become directly accessible to solution-phase, antibody-based platforms such as proximity extension assay. The extent of this leakage would be expected to track pre-analytical handling and cohort-specific collection protocols as much as underlying disease biology, a consideration relevant to any multi-site discovery cohort. This mechanism does not argue against an EV-packaged origin for CALCOCO1; rather, it suggests that the apparently free pool captured by solution-phase platforms is itself partly a function of how much vesicular cargo has already escaped by the time a sample is assayed, reinforcing the need for EV-inclusive methods when interpreting plasma proteomic findings across studies.

Conversely, proteins on the negative log2 fold-change side of the volcano plot reveal a coordinated multi-protein decrease in abundance. This depleted profile is characterized by five distinct, highly synchronized functional networks:

**1. Proliferative arrest and endosomal trafficking stagnation.** A major, highly synchronized proteomic shift involved reduced abundance and a pronounced increase in non-detection of the TBC1D3 family (including TBC1D3D, TBC1D3I, and TBC1D3E), together with TBC1D10C, in the High group. Because the TBC1D3 gene cluster has been evolutionarily specialized in driving neural progenitor proliferation and expanding cortical surface area,^49, 50^ its robust suppression suggests an early block in the brain’s baseline regenerative and neurogenic capacity. Beyond proliferation, these proteins act as specialized Rab-GTPase-activating proteins (Rab-GAPs) governing endosome-to-plasma membrane recycling and intracellular transport logistics. In healthy aged individuals (∼74 years old), maintaining rapid endosomal recycling is a vital defensive mechanism used to clear toxic extracellular amyloid-β and dynamically recycle synaptic receptors. The collective depletion of this regulatory family points to a substantial disruption of endolysosomal trafficking. This logistics failure would be expected to prevent the clearance of toxic aggregates and is consistent with upstream causal genetic variants such as BIN1, PICALM, and CD2AP identified in AD genome-wide association studies (GWAS) that establishes vesicle and endosomal trafficking failure as an early pathogenic feature of AD.^51–54^
**2. The calprotectin complex.** This module is marked by a pronounced peripheral depletion of S100A8 and S100A9, two tightly linked innate immune alarmins that form the heterodimeric calprotectin complex. Rather than a resolution of neuroinflammation, this abrupt peripheral drop is consistent with, but does not prove, exhaustive consumption and localized sequestration of circulating alarmins into central parenchymal Aβ plaques, where they may act as physical co-factors accelerating early amyloid nucleation and inflammatory mobilization.^55–57^ In the early-to-moderate stages of AD, these proteins are heavily expressed by reactive microglia surrounding diffuse plaques. Because of their ultra-high affinity for Aβ aggregates, circulating S100A8 and S100A9 are progressively drawn out of peripheral circulation and sequestered (trapped) within central parenchymal Aβ plaques, which act as a physical “sink.”^58–60^ This localized trapping causes a sharp reduction in their free, measurable concentration in plasma.
**3. The erythroid-cytoskeletal network.** We observed a broad, coordinated reduction in red blood cell membrane structural components,^61^ including spectrins (SPTA1, SPTB, SPTBN1),^62–64^ ankyrin-1 (ANK1),^65, 66^ erythrocyte membrane proteins (EPB41, EPB42),^67^ and hemoglobin subunits (HBA1, HBD).^68, 69^ This structural depletion is compatible with acquired erythrocyte membrane fragility. One possible, currently untested, contributor is mechanical shear stress and microvascular microtrauma experienced by erythrocytes when circulating through a cerebral vascular lining compromised by early cerebral amyloid angiopathy and diffuse Aβ deposition.
**4. Synaptic regression and disconnection.** This trafficking bottleneck manifests phenotypically as structural regression, evidenced by the significant decrease of Semaphorin-3C (SEMA3C).^70, 71^ Although classically studied in axon guidance during development, semaphorins are crucial for maintaining synaptic connectivity, structural plasticity, and dendritic arborization in the adult brain. The reduced detection of circulating SEMA3C in the moderate-high AD cohort may be consistent with early synaptic vulnerability; however, increased non-detection of a peripheral protein cannot by itself confirm structural dismantling of neural networks. Imaging, CSF, or tissue-level evidence would be needed to test this directly.
**5. Erasure of epigenetic and translational control.** Finally, ARID4A and NAA15 show patterns of reduced detection and decreased abundance, potentially implicating disease-associated disruption of epigenetic regulation and translational homeostasis. ^72–74^ ARID4A regulates chromatin remodeling and gene silencing architectures,^75^ while NAA15 serves as the auxiliary subunit of the NatA complex, directing co-translational N-terminal protein acetylation at the ribosome.^76, 77^ We note only as a speculative parallel, without any viral marker data in this study to support it, that the reduction of these elements superficially resembles epigenetic and translational-control changes reported during neurotropic viral reactivation (e.g., Herpes Simplex Virus 1 [HSV-1]); this analogy is not tested here and should not be read as evidence of viral involvement in this cohort.^78–80^

Paralleling this structural dissolution, our proteomic data reveal a distinct signature of metabolic collapse and enzymatic exhaustion that marks the transition into moderate-high AD. This neurodegenerative progression is highlighted by a systemic depletion in hepatic triacylglycerol lipase (LIPC) and GALNT16. The depletion of LIPC is compatible with altered peripheral-central lipid transport, consistent with a broader view of AD as having systemic metabolic components, though a single depleted protein cannot establish this on its own.^81^ Concurrently, the loss of GALNT16, which directs polypeptide O-linked glycosylation to control intracellular protein processing and secretory trafficking,^82–84^ takes on particular significance given the recent evidence implicating aberrant hypoglycosylation, or loss of protective glycan shields, as a primary post-translational driver of protein instability and aggregation in AD.^85^ The depletion of these protective enzymatic systems suggests a multi-axis failure in which defective lipid transport and compromised post-translational regulation converge to promote harmful buildup of damaged proteins inside cells.

In healthy aged individuals, balanced LIPC activity and robust GALNT16 expression maintain rigorous translational quality control to suppress protein misfolding, preserve energetic loops, and sustain non-destructive immune surveillance. During the initial transition into moderate-high AD, our data show depletion of these protective factors, consistent with (but not direct proof of) exhaustion of this defense architecture. As soluble Aβ oligomers and early tau protofibrils accumulates, the capacity of these protective enzymes and clearance matrix metalloproteinases, including MMP19, is overwhelmed. One possible, untested explanation is that these clearing enzymes are progressively depleted from systemic circulation because they become sequestered, sterically deactivated, or consumed within accumulating peripheral-central proteotoxic aggregates; direct evidence for this mechanism is not available from the present dataset.^86, 87^

Together, our identified multi-axis breakdown uncovers that early-stage AD operates as a systemic failure, rendering the peripheral-central axis highly vulnerable to subsequent metabolic collapse and terminal neurodegeneration.

Notably, external support for part of this depletion signature comes from an independent plasma proteomic study of amyloid pathology in African cohorts (the Nigerian VALIANT study, replicated in a Tanzanian cohort, and compared against the Canadian TRIAD cohort), which reported significantly lower plasma VGF in amyloid-positive relative to amyloid-negative individuals using an orthogonal, antibody-based proteomic platform (NULISA).^88^ This cross-cohort, cross-platform concordance in the direction of VGF depletion provides independent support for this specific protein-level finding; we note, however, that this replication is limited to a single protein and does not extend to the actual-causality-nominated candidates or to CALCOCO1.

Amidst these non-linear network alterations, we observed that the adhesion G protein-coupled receptor G6 (ADGRG6) was the only protein among the 3,176 quantified for which plasma abundance showed significant, moderate positive monotonic association with absolute CL values across the full clinical continuum (Spearman’s ρ = 0.49, FDR = 0.006). (**Figure 4C**). The positive association between ADGRG6 abundance and Centiloid values suggests that ADGRG6 abundance tends to increase with greater amyloid burden. This pattern is consistent with ADGRG6’s known roles in peripheral myelination, blood-brain barrier structural integrity, and vascular tissue remodeling. Within the evolving AD landscape, circulating ADGRG6 likely functions as a continuous peripheral reporter of advancing central structural damage and microvascular disruption.^89–91^ By acting as a quantitative chronometer of total amyloid burden, ADGRG6 provides a crucial biological counter-weight to the abrupt, threshold-dependent kinetics of the moderate-to-high “switch” proteins, indicating that the peripheral proteome simultaneously logs both acute homeostatic exhaustion and gradual, cumulative neurodegenerative progression.

### Actual-causality analysis prioritizes upstream drivers of the AD proteomic transition

Although differential abundance analysis has demonstrated practical utility in identifying circulating biomarkers for AD detection,^92, 93^ abundance changes alone cannot determine whether a protein lies upstream of disease progression, reflects downstream tissue injury, or simply changes with Aβ burden. This distinction is essential in AD, because therapeutic decision-making requires separating proteins that report disease stage from those that may represent therapeutic opportunities. To address this problem, we applied an actual-causality framework based on the formalism of Halpern and Pearl.^20^ Rather than claiming biological causality, this framework identifies protein assignments that satisfy increasingly rigorous empirical criteria across disease-state comparisons. Thus, differential abundance and causality analysis provide complementary information: volcano plots identify proteins that change in abundance with disease state, whereas actual-causality analysis prioritizes candidates that sit upstream of those changes.

The actual-causality (AC) system prioritizes protein assignments that exhibit causal-like behavior under increasingly rigorous empirical comparisons [see **supporting information (SI)** for details] across three primary conditions: 1) AC1 (Existence): the protein value (e.g., “high”) and the disease effect must co-occur within the same patient; 2) AC2 (Necessity and Sufficiency): approximated via “counterfactual witnesses,” where an AD-positive patient is matched to an AD-negative control who shares identical “context” features but lacks the specific protein value; and 3) AC3 (Minimality): testing singleton protein assignments to ensure the identified cause is not redundant.

Continuous protein data were binned into low, moderate, or high categories based on the 0.33 and 0.67 quantiles. Disease effect was defined by Aβ-stage severity according to CL. Candidate causes were prioritized by their uniqueness to effect cases and by extent to which their rate in affected patients exceeds their frequency in controls. Because exact matching across thousands of proteins is not feasible with high dimensional proteomic data, the analysis used context windows (C_K_) containing the top *K* ranked features. Robustness was quantified by *K_max_*, defined as the largest number of features that could be held fixed while still identifying a matched control. A higher *K_max_* indicates a candidate that remains identifiable under more stringent matching and is therefore more resistant to contextual confounding.

AC analysis identified five proteins that exhibited the highest robustness scores (**Figure 6A**). These candidates remained identifiable under stringent matching conditions requiring high feature-matching density between affected and control groups, suggesting lower sensitivity to contextual confounding. The selected proteins comprise 1) Collagen α-2(VI) chain (COL6A2; P12110), 2) FAD-dependent oxidoreductase domain-containing protein 2 (FOXRED2; Q8IWF2), 3) prolyl 3-hydroxylase 1 (P3H1; Q32P28), 4) proline-rich protein 4 (PRR4; Q16378), and 5) Golgin subfamily A member 5 (GOLGA5; Q8TBA6).

**Figure 6.**
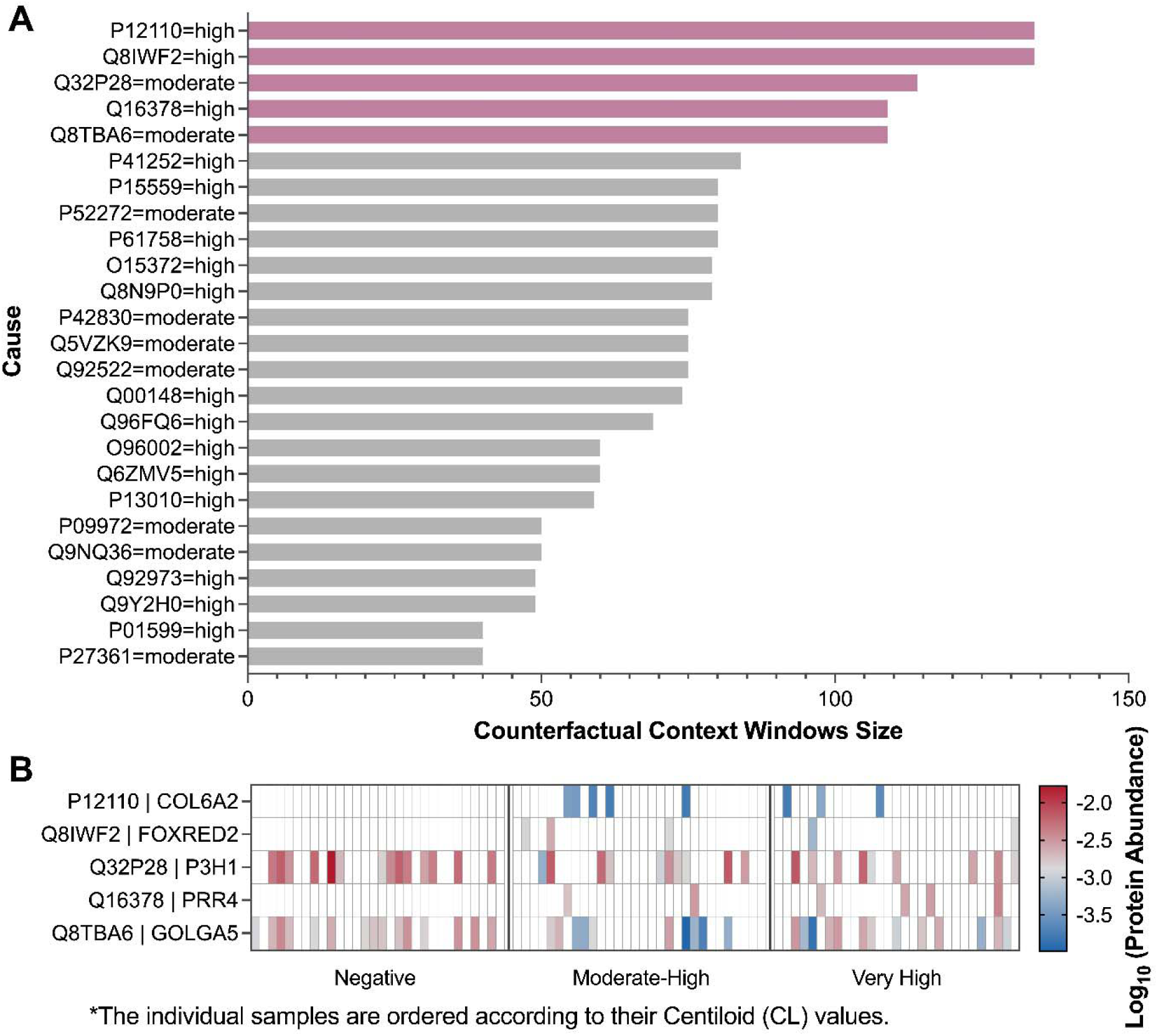
Identification of robust causal AD-associated protein signatures using Actual Causality analysis. **A)** Robustness scores for candidate protein-value causes identified by the Actual Causality (AC) framework. Each bar represents the maximum counterfactual context window size, K_max_, for a candidate protein-state association. The y-axis lists each protein candidate by UniProt ID and its AC- assigned abundance state, high or moderate. Higher K_max_ values indicate that the causal association persisted under increasingly stringent counterfactual matching conditions, requiring greater feature similarity between AD-positive and control samples, and therefore suggest greater robustness to contextual confounding. The top five ranked candidates, highlighted in purple, represent the most robust upstream causal protein signatures: COL6A2, FOXRED2, P3H1, PRR4, and GOLGA5. Gray bars indicate additional candidate causes with lower K_max_ values. **B)** a heatmap of log_₁₀_ normalized abundance for the top five prioritized causal proteins across individual samples stratified by amyloid-burden category: Aβ- negative controls, CL < 15, n = 30; moderate-to-high Aβ burden, 26 ≤ CL < 100, n = 30; and very-high Aβ burden, CL > 100, n = 30. Individual samples are ordered within each clinical group according to their Centiloid values. The heatmap illustrates stage-associated emergence or differential abundance of the top-ranked causal candidates across AD progression, with COL6A2, PRR4, and FOXRED2 showing disease-associated emergence patterns and GOLGA5 and P3H1 showing differential abundance across amyloid-burden groups.

These markers circulate at extremely low systemic levels. According to the Human Protein Atlas (HPA), only COL6A2, PRR4 and P3H1 have established plasma concentrations, with reported levels of 8.7 ng/ml, 2.4 ng/mL, and 16 pg/mL, respectively. The remaining two are primarily non-secreted intracellular proteins and lack a standardized physiological baseline in healthy blood. However, targeted research indicates that their circulating fragments typically reside in the low picogram-to- nanogram range; for instance, healthy baseline concentrations for circulating Collagen VI fragments are approximately 0.3 ng/mL in human serum.^94, 95^ Their detection and quantification are therefore consistent with the capacity of the small-molecule protein corona platform to enrich low- and ultra-low-abundance proteins and expand the measurable depth of the plasma proteome.^96^

An important consideration in interpreting the AC results is that these causal proteins are not expected to be uniformly altered in every AD-positive plasma sample (**Figure 6B**). AC is not a correlation test, rather, it identifies protein-state assignments that satisfy AC1–AC3 under matched counterfactual conditions, a substantially higher bar than detecting a cohort-level difference in abundance. Consequently, a protein can be causally informative even if it defines only a subset of patients or a particular disease stage within the broader AD-positive cohort.

When visualized as a heatmap of log_10_ mean normalized abundance across individual patient samples stratified by clinical category, these primary causal candidates exhibit distinct, stage-specific expression profiles (**Figure 6B**). Specifically, COL6A2, PRR4, and FOXRED2 show disease-stage-associated emergence patterns, whereas GOLGA5 and P3H1 were identified as robust stage-associated causes in the actual causality analysis. This coordinated shifting confirms that these prioritized candidates track with progressive amyloid accumulation. Moreover, because these candidates circulate at ultra-low blood concentrations, with baseline blood levels for COL6A2, PRR4, and P3H1, stochastic or incomplete detection across all clinical samples remains a technical limitation for conventional plasma proteomic platforms. Consequently, validating their clinical utility will require highly sensitive, targeted quantification frameworks, such as enzyme-linked immunosorbent assays (ELISA) or alternative ultra-sensitive immunoassays engineered for the low-abundance sub-proteome.

### Biological significance and disease-related links

The identified AC candidates highlight a multifaceted breakdown of cellular homeostasis in AD, spanning from peripheral extracellular matrix (ECM) structural failure to central organelle degradation such as Golgi fragmentation.

First, Collagen α2(VI), encoded by COL6A2, is a key subunit of the Collagen VI complex, which preserves ECM structural integrity. Independent support for this ECM-centered signature comes from a large prospective proteomic study (UK Biobank, n = 52,645, 14.1-year follow-up), in which genome-wide enrichment analysis of dementia-associated plasma proteins identified ECM organization and collagen-containing extracellular matrix among the top enriched pathways, consistent with a broader role for ECM disruption in dementia risk beyond the specific proteins nominated here.^97^ In addition to this structural role, Collagen VI is neuroprotective^98^ where neurons upregulate its production in response to Aβ,^98^ partly by sterically blocking interactions between Aβ oligomers and the neuronal surface.^99^ By forming a pericellular network, Collagen VI physically impedes Aβ binding to neurons, thereby preventing downstream toxicity and cell death.^98, 100^ Collagen VI also promotes aggregation of soluble Aβ into larger, less toxic assemblies,^98, 101^ reducing their capacity to damage the vulnerable cell membrane.

Second, FAD-dependent oxidoreductase domain-containing protein 2 (FOXRED2) is upregulated by Aβ in AD.^102^ Elevated FOXRED2 impairs proteasome function, leading to accumulation of damaged and misfolded proteins and ultimately promoting neuronal death.^102^

Third, P3H1 is a vital enzyme that, alongside CRTAP and Cyclophilin B, catalyzes the post-translational hydroxylation of proline residues. This modification is essential for properly folding, stabilizing, and assembling collagen within the endoplasmic reticulum.^103–105^

Fourth, proline-rich protein 4 (PRR4), primarily identified in tear-fluid proteomics, has been recently associated with AD progression, suggesting its potential as a noninvasive biomarker.^106–109^

Fifth, GOLGA5 (Golgin subfamily A member 5) is a Golgi structural protein involved in maintaining Golgi architecture and vesicle trafficking. Golgi fragmentation is an early and prominent feature of AD.^110, 111^ The fragmentation of the Golgi apparatus creates a toxic feedback loop in AD: the loss or structural modification of vital structural proteins like GOLGA5 (Golgin-84) impairs intracellular transport, which accelerates the clearance failure of amyloid precursor protein processing products and triggers tau hyperphosphorylation.^112, 113^

### Mechanistic convergence: connecting actual causality to volcano plot phenotypes

The five upstream causal candidates prioritized by our AC framework are not isolated anomalies; rather, their distinct, stage-specific expression profiles provide a mechanistic bridge to the multi-systemic network collapse observed in the mid-stage volcano plot “switch” phenotypes. By mapping their individual biochemical functions onto the broader proteomic landscape, we uncover an integrated network of homeostatic failures that links progressive amyloid burden to peripheral-central tissue dissolution.

#### I. Organelle stagnation and intracellular trafficking failure (GOLGA5/FOXRED2 CALCOCO1/TBC1D3)

The upstream prioritization of GOLGA5 by the actual causality analysis may help contextualize the endolysosomal trafficking bottleneck reflected by proteins with negative log2 fold changes in the volcano plot. When GOLGA5-mediated Golgi architecture fragments, vesicular trafficking logistics between the endoplasmic reticulum, Golgi, and plasma membrane are paralyzed. This transport failure may be further compounded by the disease-associated emergence of FOXRED2, a candidate causal driver with reported links to impaired proteasome function in other systems. If a similar effect occurs here, a fragmented Golgi and reduced proteasome capacity could plausibly limit clearance of misfolded proteins and trapped cargo. This is one possible, not directly tested, explanation for the high-magnitude accumulation of CALCOCO1 in the mid-stage volcano plot; the cell is forced to aggressively upregulate this reticulophagy cargo receptor as an emergency, auxiliary stress response to clear the expanding endoplasmic reticulum sub-compartment trafficking issues. Ultimately, this structural bottleneck coincides with a marked loss of detectable TBC1D3 Rab-GAP family proteins; while consistent with upstream organelle fragmentation contributing to the endosomal recycling collapse seen at the mid-stage transition, this cross-sectional proteomic data cannot establish that one directly triggers the other.

#### II. Extracellular matrix disruption and neurovascular microtrauma (P3H1/COL6A2 Spectrins/ANK1/SEMA3C)

A second axis of convergence links the causal matrix-modifier candidates to the structural collapse of the erythroid-cytoskeletal network. P3H1 is natively required for post-translational hydroxylation and stable assembly of Collagen Type IV while COL6A2 co-deposits at active cerebral amyloid angiopathy fronts. The progressive dysregulation and depletion of P3H1 yields an unstable, structurally compromised vascular basement membrane that breaches BBB integrity. As the vascular lining degrades due to defective collagen folding and active Aβ deposition, circulating red blood cells may experience greater mechanical shear stress and microvascular friction. This vascular microtrauma is one plausible, untested contributor to the sweeping, coordinated reduction of membrane structural components, namely -spectrin (SPTA1), -spectrin (SPTB), and ankyrin-1 (ANK1), on the depleted side of the volcano plot. Furthermore, this structural and neurovascular matrix disintegration may plausibly extend to the parenchymal microenvironment; the acute peripheral depletion of the synaptic plasticity anchor SEMA3C is consistent with this picture, although it does not by itself establish physical dismantling of neural networks.

#### III. Systemic epithelial barrier and exocrine failure (PRR4 S100A8/S100A9)

Finally, the actual causality ranking of PRR4 suggests that early AD pathobiology is an organismal-wide phenomenon rooted in shared autonomic and barrier regulatory circuits. PRR4 dysregulation mirrors systemic epithelial and exocrine barrier vulnerability. As these mucosal and neurovascular boundaries are compromised, a coordinated systemic alarmin response may be triggered. This provides a plausible link to the marked reduction in the calprotectin complex (S100A8/S100A9) observed during the mid-stage transition. Rather than a simple downregulation of production, the initial barrier stress and advancing amyloid deposition may also indicate an active peripheral vacuum or “sink effect,” drawing free circulating S100A8/A9 alarmins out of the plasma and trapping them into central parenchymal Aβ plaques where they act as direct binding co-factors. Separately, we note that TNFSF10 (TRAIL), which we found depleted in the moderate-to-high cohort (**Figure 3A** and **B**), is the ligand for TNFRSF10B, a death receptor independently identified as significantly associated with incident all-cause dementia risk in the same large prospective UK Biobank study; while this cross-cohort observation involves the ligand and receptor rather than the same protein, it is consistent with altered TRAIL/death-receptor signaling as a feature of dementia risk more broadly.^97^ In addition, consistent with our findings, a recent large-scale plasma proteomic study of neurodegenerative dementias shows TNFSF10 decreased across various stages of AD as compared to the control healthy plasmas.^43^

### Deep plasma proteomics enables therapeutic staging of AD

This study demonstrates that deep, small-molecule-modulated protein corona plasma proteomics combined with counterfactual causal inference can resolve stage-associated molecular transitions along the AD continuum. Our findings support a model in which AD progression cannot be interpreted or managed through absolute Aβ burden alone. Rather than progressing as a gradual, uniform continuum, the peripheral proteome undergoes a pronounced, biological “switch” during the moderate-to-high amyloid transition. This window is characterized by high-magnitude accumulation of the reticulophagy cargo receptor CALCOCO1 alongside a multi-protein systemic depletion of the S100A8/S100A9 calprotectin complex, core erythroid-cytoskeletal elements (SPTA1, SPTB, ANK1), and the TBC1D3 endosomal trafficking family. Crucially, this threshold-driven switch is complemented by the continuous, linear escalation of ADGRG6, which serves as a progressive quantitative tracking anchor for absolute disease burden across the entire spectrum. This entire peripheral-central state goes back to the negative levels in advanced disease (CL > 100), defining a distinct, blood-detectable transition from early homeostatic compensation to structural exhaustion.

If replicated and validated against clinical outcomes, this staging framework could eventually inform discussions of disease-modifying therapies. We emphasize that this is a hypothesis generated from a single cross-sectional cohort of 90 individuals and has not been tested against treatment response; it should not presently be used to guide therapeutic decisions. Prospective, longitudinal studies with treatment-outcome data would be required before any claim that low-abundance proteome signals predict reduced benefit from anti-Aβ immunotherapies, or that individuals below this proteomic threshold preferentially benefit from early Aβ-focused interventions, could be supported. These findings align with the proposed “turning point” in the cellular phase of AD described by De Strooper and Karran,^1^ and extend that concept by mapping the peripheral window where homeostatic defenses fail before irreversible, widespread neurodegeneration takes hold.

## Discussion

Overall, our platform successfully distinguishes correlation-derived staging biomarkers from actual-causality-prioritized candidates, including the disease-associated emergence of COL6A2, PRR4, and FOXRED2, alongside stage-associated abundance changes in GOLGA5 and P3H1.This provides a dual-axis framework for clinical decision-making: matching patients to therapies according to the active molecular state of their disease, and selecting drugs based on the specific biological processes driving progression at that stage. To transition these trace discoveries into scalable clinical tools, future studies should deploy targeted, ultra-sensitive multiplex immunoassays optimized to track the precise stoichiometric ratio of CALCOCO1 accumulation and the positive association of ADGRG6 with CL burden, and the concurrent depletion of selected proteins, thereby enabling more rigorous evaluation of stage-specific biomarker signatures in clinical trials.

Beyond the specific proteins nominated here, this study’s central contribution is methodological: separating markers that are merely correlated with disease stage from those whose behavior is most consistent with a causal role in the transition between stages. Most large-scale AD plasma proteomic efforts to date, including recent population-scale studies of incident dementia risk,^&#x25A1;&#x25A1;.,&#x25A1;¹^ have relied on differential-abundance or hazard-ratio associations to nominate candidate biomarkers. These approaches are well suited to answering whether a protein’s level differs between groups or predicts a future binary outcome, but they cannot, by construction, distinguish a protein that changes because it is swept along by disease progression from one that actively participates in driving that progression. By applying an actual-causality framework^²&#x25A1;^ alongside conventional differential-abundance testing, we could ask a different, complementary question: among the proteins that change, which are most consistent with occupying an upstream, mechanistically load-bearing position in the transition itself. This distinction matters for translational prioritization, because it is the causally-implicated proteins, not necessarily the most statistically significant ones, that are the more defensible starting points for target validation and functional follow-up.

Placed alongside the broader blood-based AD biomarker literature, our findings occupy a different, complementary niche. Large prospective cohort studies such as the UK Biobank proteomic analysis of incident dementia^&#x25A1;¹^ and large-scale case-control plasma proteomic profiling efforts have established GFAP, NEFL, GDF15, and related markers as robust predictors of the transition from cognitively normal to clinically diagnosed dementia over years to decades. These studies answer a prediction question: who, among currently unimpaired individuals, will progress to dementia. Our study instead asks a staging question within a population already positioned along the amyloid continuum: among individuals who are already Aβ-positive, does the peripheral proteome distinguish those in an early, potentially more therapeutically responsive state from those in a later, more entrenched one. These are related but distinct clinical problems, and we see our platform as complementary to, rather than competing with, established risk-prediction biomarkers; a combined framework that first identifies at-risk individuals using established predictors and then stages their active molecular biology using an approach such as ours could in principle support both earlier identification and more precise therapeutic sequencing, though this combination has not been tested here and would require dedicated prospective study.

Some convergence with independent cohorts is worth highlighting, with appropriate caution. VGF, one of the proteins depleted in our moderate-to-high cohort, was independently reported as significantly reduced in amyloid-positive relative to amyloid-negative individuals in the VALIANT and TRIAD cohorts spanning Nigerian, Tanzanian, and Canadian populations using an orthogonal antibody-based platform.^&#x25A1;²^ Because this convergence involves a single protein identified through differential abundance rather than our causal-inference pipeline, it should be read as modest external support for the plausibility of our differential-expression findings specifically, not as validation of the actual-causality-nominated candidates, which remain unreplicated in any independent cohort.

Mechanistically, the coordinated nature of the changes marking the moderate-to-high transition, spanning autophagy receptor accumulation, calprotectin depletion, erythroid- cytoskeletal loss, and endosomal trafficking collapse, is more consistent with a systemic proteostatic and vesicular-trafficking failure than with an isolated single-pathway defect. This pattern is compatible with, though it does not on its own establish, the broader model in which AD transitions through a limited number of qualitatively distinct cellular phases rather than accumulating damage in a purely linear fashion, a view articulated in the proposed cellular phase and turning-point frameworks for AD.^¹·²^ Our data extend this conceptual model by suggesting that such a phase transition, if it exists, may leave a coordinated, multi-protein signature detectable in peripheral blood rather than only in brain tissue or cerebrospinal fluid, which would be of practical value given the relative inaccessibility of the latter two compartments for repeated sampling.

The behavior of ADGRG6 offers a useful counterpoint to the discrete-switch proteins. Rather than emerging abruptly at a single transition, ADGRG6 tracked amyloid burden continuously across the full range studied. Adhesion G protein-coupled receptors of this family have established roles in myelination, vascular development, and maintenance of blood–brain barrier integrity,^&#x25A1;^3^_&#x25A1;&#x25A1;^ processes plausibly relevant to progressive, cumulative vascular and structural change rather than to a discrete cellular-state transition. If this distinction holds under future validation, it would suggest that a clinically useful staging panel may need to combine at least two qualitatively different classes of markers: discrete switch-like proteins that flag whether a threshold has been crossed, and continuous tracking proteins that quantify graded burden independent of that threshold, in the same way that clinical staging in other chronic diseases often combines categorical and continuous measures rather than relying on either alone.

These findings, taken together, motivate a two-track path toward clinical translation. The first track is analytical: the candidate proteins identified here, particularly the actual- causality-nominated set, circulate at concentrations well below the sensitivity of most routine clinical assays, and translating them into a usable tool will require targeted, ultra-sensitive, multiplexed immunoassays rather than untargeted discovery proteomics. The second track is clinical: any eventual staging panel would need to be evaluated not only for its association with amyloid burden but for its ability to stratify patients by treatment response in prospective interventional cohorts, since the ultimate clinical value of a staging biomarker lies in whether it changes therapeutic decisions rather than in its statistical association with existing disease-severity measures. Neither track is complete here; we regard these findings as hypothesis-generating groundwork for both, not a validated clinical tool.

Finally, we note that the dual-axis framework distinguishing correlational stage markers from causally-implicated drivers may have relevance beyond Alzheimer’s disease. Many chronic, slowly progressive conditions are currently staged primarily by cumulative burden of a single pathological hallmark, analogous to amyloid burden in AD, even though the clinical trajectory and treatment responsiveness of individual patients at a given burden level can vary substantially. To the extent that our approach, combining deep low-abundance plasma proteomics with a causal-inference layer, generalizes to other conditions with a similar mismatch between cumulative pathological burden and active disease state, it may offer a template for staging frameworks beyond the specific proteins and disease reported here. We offer this as a direction for future methodological work rather than as an established generalization, since it has not been tested in any condition other than AD in this study.

It is worth situating the actual-causality approach itself within the broader epistemics of observational biomarker discovery. Actual causality, as formalized by Halpern and Pearl,^²^ provides a principled way to ask which observed variables are most consistent with occupying a causally load-bearing position given a specified structural model and the observed data, but it remains fundamentally a inference over cross-sectional, observational measurements rather than a substitute for interventional evidence. We view its value here as one of prioritization rather than proof: among thousands of candidate proteins, it offers a principled, pre-specified basis for allocating scarce downstream validation resources, such as targeted assay development and functional perturbation studies, toward a tractable shortlist, rather than asserting that the nominated candidates are confirmed causal drivers of disease progression. This framing is consistent with how causal-inference methods are increasingly used across observational biomedicine: as a rigorous filter narrowing an intractably large hypothesis space, not a replacement for experimental validation.

### Limitations

We acknowledge several limitations. First, the cohort (n = 30 per group; N = 90) is modest compared with the number of proteins tested (3,176); larger, independently ascertained cohorts are required to validate reported associations. Second, the study lacks an independent validation cohort and orthogonal quantification (e.g., ELISA) of candidate biomarkers and putative causal drivers; such validation is essential before any identified proteins can be considered for clinical use.

## Consent statement

The AIBL study — including its follow-up protocol and all subsequent amendments and revisions — was approved by the institutional human research ethics committees of Austin Health, St Vincent’s Health, Hollywood Private Hospital, and Edith Cowan University. All participants provided written informed consent prior to undergoing study assessments, and the study was conducted in accordance with the Declaration of Helsinki.

