## Supplementary material for "Deep nanoparticle protein corona plasma proteomics resolves a stage-specific peripheral signature of Alzheimer’s disease": SI

### 1 Methodology

#### 1.1 Overview

The core objective of actual causality analysis was to identify protein values that are plausible actual causes of Alzheimer's disease (AD) in a patient-level protein dataset. The analysis follows the original Halpern and Pearl definition of actual causality[1] but implements it as an approximate search procedure using the observational data as supporting evidence.

Formally, let

$$M = (\mathcal{U}, \mathcal{V}, \mathcal{F})$$

be a structural causal model with exogenous variables $\mathcal{U}$, endogenous variables $\mathcal{V}$, and structural equations $\mathcal{F}$. Let $\vec{u}$ be the actual context, let $\varphi$ be an effect formula over variables in $\mathcal{V}$, and let $\vec{X} = \vec{x}$ be a candidate cause. In the original Halpern Pearl definition, $\vec{X} = \vec{x}$ is an actual cause of $\varphi$ in $(M, \vec{u})$ if the following conditions hold:

**AC1.** The candidate cause and the effect are both true in the actual context:

$$(M, \vec{u}) \models \vec{X} = \vec{x} \quad \text{and} \quad (M, \vec{u}) \models \varphi. \tag{AC1}$$

**AC2.** There exists a partition $(\vec{Z}, \vec{W})$ of the endogenous variables $\mathcal{V}$, with $\vec{X} \subseteq \vec{Z}$, an alternative setting $\vec{x}'$ of $\vec{X}$, and a contingency setting $\vec{w}$ of $\vec{W}$ such that, writing $\vec{z}^*$ for the actual values of $\vec{Z}$ in $(M, \vec{u})$, the following two requirements hold:

$$(M, \vec{u}) \models [\vec{X} \leftarrow \vec{x}', \vec{W} \leftarrow \vec{w}]\neg\varphi, \tag{AC2(a)}$$

and, for every subset $\vec{Z}' \subseteq \vec{Z}$,

$$(M, \vec{u}) \models [\vec{X} \leftarrow \vec{x}, \vec{W} \leftarrow \vec{w}, \vec{Z}' \leftarrow \vec{z}^*]\varphi. \tag{AC2(b)}$$

AC2(a) is the necessity condition. Changing the actual cause under a suitable contingency changes the effect. AC2(b) is the sufficiency condition. When the actual cause is present, the effect remains under the same contingency used in AC2(a).

**AC3.** No strict subassignment of $\vec{X} = \vec{x}$ satisfies the preceding conditions. This excludes causes that contain irrelevant conjuncts.

In our analysis, the structural model $M$ and its equations are not estimated. Instead, the definition is used to guide an observation based analog. Each patient row is treated as an observed context, the effect formula $\varphi$ is the event that the patient's severity belongs to the selected AD severity set, and each candidate cause is a singleton protein value assignment

$$X_{j,s} \equiv (B_j = s).$$

In this analog, AC1 requires the candidate protein value and the effect to co-occur in actual effect patients. AC2(a) is approximated by searching for matched control rows that agree with an effect row on a selected context window but differ in the candidate protein value and lack the effect. The need for a context window arises from the large number of features available, making a complete match of non-causal features unlikely given the size of the dataset. AC2(b) is approximated by observing features in effect rows, then selecting context features and treating those features as the contingency variables in $\vec{W}$ for the empirical comparison. Because we test singleton protein value assignments, AC3 is satisfied within the tested cause class, but we do not claim minimality over untested multi-protein value conjunctions.

Accordingly, the method should be interpreted as an actual-causality-inspired screening procedure rather than a proof of actual causality in the full biological system. A protein value is retained when it is observed in effect patients, has an empirical counterfactual witness among matched controls, and remains supported as the context window used for matching becomes more restrictive.

#### 1.2 Data Representation

Let the dataset contain $n$ patients and $p$ measured proteins. For patient $i$, let

$$S_i \in \{0, 1, 2, 3\}$$

denote disease severity, with 0 representing negative disease status and larger values representing increasing severity. A specified set of severity levels $\mathcal{L}$ defines the effect of interest:

$$Y_i = \mathbf{1}\{S_i \in \mathcal{L}\}.$$

For our analysis our effect is defined as:

$$\varphi := \big(Y_i = \mathbf{1}\{S_i \in \{1, 2, 3\}\}\big)$$

Patients with $Y_i = 1$ are treated as effect cases, and patients with $Y_i = 0$ are treated as controls. This allows the testing of different effects, such as any disease severity, moderate disease only, high disease only, or very high disease only.

Raw protein measurements are continuous. Since the actual-causality screen operates on discrete variable assignments, each protein $j$ is converted into a categorical variable

$$B_{ij} = b_j(R_{ij}),$$

where $R_{ij}$ is the raw measurement and $b_j$ is a configurable binning function. Each protein is assigned to low, moderate, or high according to protein-specific lower and upper quantiles. For this analysis we selected the lower and upper quantiles to be 0.33 and 0.67 respectively. Missing values are treated as zero before binning, matching the assumption that blank measurements correspond to no detected protein abundance.

#### 1.3 Candidate Causes

A candidate cause is a protein value assignment

$$X_{j,s} \equiv (B_j = s),$$

where $j$ indexes a protein and $s$ is one of its binned ranges. For example, a candidate may be "protein $x$ is high." For each candidate, we count how often the value appears in effect patients and control patients:

$$n_E(j, s) = \sum_i \mathbf{1}\{Y_i = 1, B_{ij} = s\},$$

$$n_C(j, s) = \sum_i \mathbf{1}\{Y_i = 0, B_{ij} = s\}.$$

For our analysis, in order to satisfy AC2, we only consider candidate causes which are unique to the rows where the effect occurs and exclude those that appear in control rows.

##### 1.3.1 Feature Ranking for Context Selection

Based on $n_E(j, s)$ and $n_C(j, s)$, their corresponding empirical rates are

$$p_E(j, s) = \Pr(B_j = s \mid Y = 1), \qquad p_C(j, s) = \Pr(B_j = s \mid Y = 0).$$

The candidate table is sorted to prioritize protein values that are most causally active in the observed data. The ranking first prioritizes values that are unique to effect cases, then values with larger rate differences

$$\Delta(j, s) = p_E(j, s) - p_C(j, s),$$

and then values that occur in more effect patients.

This ranked table solely used as a way of identifying a tractable subset of protein value candidates worth testing against the actual-causality conditions, and is not used directly in the causal analysis.

#### 1.4 Satisfying AC Requirements

### 1.4.1 AC1

In our analysis, AC1 is evaluated at the patient-row level. A patient $i$ supports candidate $X_{j,s}$ if

$$B_{ij} = s \wedge Y_i = 1.$$

Due to the method in which we identify candidates from the observational data, we only test protein values observed in rows where the effect is present. Thus, AC1 is satisfied for all proposed candidates.

### 1.4.2 AC2(a)

AC2(a) requires evidence that changing the candidate cause can change the effect under some contingency. As the dataset is observational, we approximate this counterfactual condition using matched control rows. Let $C$ denote a selected context window, i.e., a subset of protein features that will be held fixed during matching. For a candidate $X_{j,s}$, the tested protein $j$ is removed from the matching context so that the context is

$$C_{-j} = C \setminus \{j\}.$$

An effect patient $i$ and a control patient $k$ form a counterfactual witness for $X_{j,s}$ if

$$Y_i = 1, \quad B_{ij} = s,$$

$$Y_k = 0, \quad B_{kj} \neq s,$$

and

$$B_{i\ell} = B_{k\ell} \quad \text{for every } \ell \in C_{-j}.$$

In words, the effect patient has the candidate cause and the effect, the matched control patient lacks the candidate cause and lacks the effect, and all selected context features are identical between them. This is an analog of the intervention in AC2(a): under a fixed context, the candidate feature differs and the effect differs.

The candidate satisfies AC2(a) if at least one supporting effect row has such a matched control witness.

### 1.4.3 AC2(b)

AC2(b) requires that the effect remain stable when the candidate cause is held at its actual value and relevant background variables are fixed. In our approach, we only consider candidates whose values are unique to effect rows, so AC2(b) is approximated with respect to our observational data.

We approximate AC2(b) by matching on the values of the selected context variables. The context window represents the part of the causal model that is being treated as fixed. Formally, the matched comparison keeps

$$B_{iC_{-j}} = B_{kC_{-j}},$$

so the observed difference between the effect row and the control row is only in the candidate protein's value.

This should be interpreted as an empirical AC2(b)-style screen rather than a proof of actual causality in the underlying model that our data represents. The quality of the approximation depends on whether the selected context window contains the relevant causal variables and whether matched control rows exist.

#### 1.5 Feature Subset Construction

The proteomics dataset contains thousands of protein variables (see Figure 1 of the main manuscript and Supplementary Dataset). Exact matching on all variables would be the closest observed analog of a complete causal model, but in high-dimensional data this is usually impractical. Most patient rows become unique once enough protein values are held fixed. As a result, a full-row context often produces few or no counterfactual matches.

To address this, we construct smaller models from subsets of the most causally active features. The ranking described above identifies proteins whose values are most enriched in the effect group. Context windows are then selected from this ranked set. Our context selection method requires that the top $K$ ranked proteins are matched, where $K$ is configurable. This method provides a controlled way to test over increasing context sizes. Small windows give more matched controls but represent a coarser causal model. Larger windows require more variables to be equal between patients and therefore more closely approximate the full observed model.

#### 1.6 Context Windows and Robustness

Let $C_K$ denote a context window containing the top $K$ ranked features. For each candidate cause, we then record the largest context size for which the cause survives counterfactual screening:

$$K_{\max}(j, s) = \max\{K : X_{j,s} \text{ survives matching under } C_K\}.$$

This value is used as a robustness measure. A larger value of $K_{\max}$ means that the candidate remained valid even when a larger number of protein features had to match exactly when searching for a supporting counterfactual. Thus, larger context windows imply comparisons that are closer to the actual high-dimensional model.

#### 1.7 Cause Interpretation

Our methodology identifies protein value assignments that satisfy empirical analogs of AC1 and AC2 under selected causal submodels. AC1 is satisfied when the candidate value and the disease effect are jointly observed in actual patient rows. AC2(a) is supported when a matched control row exists with the same context, a different candidate value, and no effect. AC2(b) is approximated by holding the selected context features fixed during matching.

Because the full biological causal model is unknown and the dataset is high-dimensional, the method does not claim to prove causality in the full observed dataset. Instead, it provides a principled procedure for finding protein values that behave like actual causes under increasingly rich empirical models. The most compelling candidates are those that are enriched in effect patients, have matched counterfactual witnesses, and remain valid as the context window expands.

### 2 Supplemental Tables

Table 1: HPLC conditions for the 24 SPD gradient method

| Gradient Specifications and LC Configuration | | | |
|---|---|---|---|
| **Gradient** | **Time (min)** | **% Mobile Phase B** | **Flow ($\mu$l/min)** |
| | 0 | 8 | 0.5 |
| | 2.5 | 8 | 0.5 |
| | 3.0 | 8 | 0.25 |
| | 37.0 | 22.5 | 0.25 |
| | 48.5 | 35 | 0.25 |
| | 48.9 | 98 | 0.25 |
| | 49.0 | 98 | 0.5 |
| | 54.0 | 98 | 0.5 |
| **LC Parameters (Trap and Elute Configuration)** | | | |
| Fast Loading/Equilibration Mode | | Pressure Control | |
| Loading/Equilibration/Wash Pressure | | Max Pressure | |
| Equilibration Factor | | 3 | |
| Sampler Temperature | | 7°C | |
| Mobile Phase A / Weak Wash | | 0.1% Formic Acid in Water | |
| Mobile Phase B / Strong Wash | | 0.1% Formic Acid in 80% Acetonitrile | |
| Zebra Wash | | Enabled | |
| Zebra Wash Cycles | | 4 | |
| Analytical Column Temperature | | 50°C | |
| **Column Specifications** | | | |
| Analytical Column | EASY-Spray PepMap Neo Column, 2 $\mu$m C18, 75 $\mu$m × 50 cm (P/N ES75500PN) | | |
| Trap Column | PepMap Neo Trap Cartridge, 5 $\mu$m C18, 300 $\mu$m × 5 mm (P/N 174500) | | |

Table 2: HPLC conditions for the 60 SPD gradient method

| **60 SPD Gradient Specifications and LC Configuration** | | | |
|---|---|---|---|
| **Gradient** | **Time (min)** | **% Mobile Phase B** | **Flow ($\mu$l/min)** |
| | 0 | 10 | 2.0 |
| | 0.3 | 10 | 2.0 |
| | 0.6 | 10 | 0.8 |
| | 13.6 | 22.5 | 0.8 |
| | 20.5 | 35.0 | 0.8 |
| | 20.9 | 55.0 | 2.0 |
| | 20.95 | 99.0 | 2.0 |
| | 22.35 | 99.0 | 2.0 |
| **LC Parameters (Trap and Elute Configuration)** | | | |
| Fast Loading/Equilibration Mode | | Pressure Control | |
| Loading/Equilibration/Wash Pressure | | Max Pressure | |
| Equilibration Factor | | 3 | |
| Sampler Temperature | | 7°C | |
| Mobile Phase A / Weak Wash | | 0.1% Formic Acid in Water | |
| Mobile Phase B / Strong Wash | | 0.1% Formic Acid in 80% Acetonitrile | |
| Zebra Wash | | Enabled | |
| Zebra Wash Cycles | | 4 | |
| Analytical Column Temperature | | 50°C | |
| **Column Specifications** | | | |
| Analytical Column | EASY-Spray PepMap Column, 2 $\mu$m C18, 150 $\mu$m × 15 cm (P/N ES906) | | |
| Trap Column | PepMap Neo Trap Cartridge, 5 $\mu$m C18, 300 $\mu$m × 5 mm (P/N 174500) | | |

Table 3: Orbitrap Astral Zoom Mass Spectrometer parameters for 60SPD and 24 SPD methods.

**(A) Global source and mass spectrometer parameters**

| **Parameter** | **Value** |
|---|---|
| Positive Ion Voltage | 2100 Volts |
| Ion Transfer Tube Temperature | 290°C |
| Expected Peak Width | 10 seconds (24SPD), 6 seconds (60SPD) |
| Default Charge State | 2 |
| Lock Mass Correction | Off |

**(B) MS1 full scan experiment parameters**

| **Parameter** | **Value** |
|---|---|
| Orbitrap Resolution | 240K |
| Scan Range ($m/z$) | 380–980 |
| RF Lens (%) | 40 |
| Normalized AGC Target (%) / Absolute AGC Value | 500% / 5.00e6 |
| Maximum Injection Time | 5 milliseconds |
| Microscans | 1 |

**(C) MS2 DIA scan experiment parameters**

| **Parameter** | **Value** |
|---|---|
| Precursor Mass Range ($m/z$) | 380–980 |
| Isolation Window ($m/z$) | 2.5 (24SPD), 3 (60SPD) |
| Window Placement Optimization | On |
| AGC Target | Custom |
| Normalized AGC Target (%) / Absolute AGC Value | 500% / 5.00e4 |
| Maximum Injection Time | 7 milliseconds |
| DIA Scan Range ($m/z$) | 150–2000 |
| HCD Collision Energy (%) | 25 |
| RF Lens (%) | 40 |
| Pre-Accumulation | On |
| Loop Control | Time |
| Time | 0.6 seconds |
